# The *Dictyostelium discoideum – Mycobacterium marinum* infection model, a powerful high throughput screening platform for anti-infective compounds

**DOI:** 10.1101/2024.12.03.626613

**Authors:** Jahn Nitschke, Nabil Hanna, Thierry Soldati

## Abstract

Tuberculosis is among the world’s deadliest diseases, causing approximately 2 million deaths annually. The urgent need for new antitubercular drugs has been intensified by the rise of drug-resistant strains. Despite recent advancements, most hits identified through traditional target-based screening exhibit limited efficacy *in vivo*. Consequently, there is a growing demand for whole-cell-based approaches that directly utilize host-pathogen systems. The *Dictyostelium discoideum*–*Mycobacterium marinum* host–pathogen system is a well-established and powerful alternative model system to study mycobacterial infections. In this article, the phenotypic host-pathogen protocol assay is presented here which relies on monitoring *M. marinum* during its infection of the amoeba *D. discoideum*. This assay is characterized by its scalability for high-throughput screening, robustness, and ease of manipulation, making it an effective system for compound screening. This system provides not only bacterial load readout via a bioluminescent *M. marinum* strain, but now also host survival and growth via a fluorescent *D. discoideum* strain enabling further host characterization by quantifying growth inhibition and potential cytotoxicity. Finally, the system was benchmarked with selected antibiotics and anti-infectives and calculated IC_50_s and MICs where applicable, demonstrating its capability to differentiate between antibiotics and anti-infective compounds.

**Importance:** This methods paper introduces a robust, scalable, and high-throughput phenotypic host-pathogen assay based on the well-established *Dictyostelium discoideum–Mycobacterium marinum* system. In contrast to conventional target-based drug screening approaches, which often struggle to translate effectively *in vivo*, this platform directly monitors pathogen-host interactions, providing comprehensive insights into bacterial load, host survival, and potential cytotoxicity. By employing bioluminescent *M. marinum* and fluorescent *D. discoideum* strains, we validated the system using established antibiotics and anti-infective compounds, effectively distinguishing their effects through IC50 and MIC calculations.

## Introduction

*Dictyostelium discoideum* (Dd) is a social amoebae and professional phagocyte. As such, it shares conserved and fundamental cell autonomous immunity strategies with animal innate immune phagocytes. Additionally, it is genetically tractable, a fully sequenced model organism (Eichinger et al., 2005), and readily infected with intracellular pathogens, such as *Legionella pneumophila* (Solomon & Isberg, 2000; Welin et al., 2023) but also *Vibrio cholerae*, *Francisella noatunensis*, *Pseudomonas aeruginosa*, *Salmonella enterica* and *Mycobacterium* species as reviewed (Dunn et al., 2018). Mycobacteria are a genus of bacteria characterized by their waxy cell wall (Batt et al., 2020) and several members of strains pathogenic for human and animals, foremost the *Mycobacterium tuberculosis* (Mtb) complex (Kanabalan et al., 2021). A close relative of the Mtb complex is *Mycobacterium marinum* (Mm) (Stinear et al., 2008), a facultative human pathogen, able to create skin lesions similar to lung lesions observed during active tuberculosis (TB) (Ramakrishnan, 2004; Tobin & Ramakrishnan, 2008). Mm is a convenient substitute for Mtb, since it conserves virulence strategies, but has a doubling time as short as 8 hours and requires infrastructure of biological safety level 2, instead of 3 (Ramakrishnan, 2004). Drugs against TB have often been identified by screens performed on attenuated strains such as the *Mycobacterium bovis* Bacillus Calmette-Guérin strain (Taneja & Tyagi, 2007), used as a live vaccine (Calmette et al., 1927) or even a non-pathogenic strain, such as *Mycobacterium smegmatis* (Altaf et al., 2010; Andries et al., 2005). While this screening strategy is resourceful, it is accompanied by a significant attrition rate due to the disregard of the complex mycobacterial infection biology. As an alternative, screening on infected host cells, such as murine Raw264.7 macrophages or human THP-1 macrophages, has become an accepted strategy (Christophe et al., 2010; Pethe et al., 2013; Schaaf et al., 2016). This is not only incorporating infection biology parameters and aspects of pharmacokinetics, but also opens the door to anti-infective drugs that target only or additionally the host (Tobin, 2015; Udinia et al., 2023). However, the bottleneck of phenotypic screens is the progression to a mode of action, which is even more difficult when working with systems that are resource-intensive to maintain, and hard to manipulate genetically, such as Mtb or human primary macrophages. Therefore, we advocate a Dd – Mm infection model to close this gap. We recently demonstrated the power of this model to investigate host pathogen interactions (López-Jiménez et al., 2018; Raykov et al., 2023), but also to detect active anti-infective compounds (Hanna et al., 2020; Kirchhoffer et al., 2023, 2024; Nitschke et al., 2024; Trofimov et al., 2018). Consequently, we developed this system to the level of a high throughput platform that monitors not only bacterial load via a bioluminescent Mm strain (Andreu et al., 2010; Arafah et al., 2013), but now also host survival and growth via a fluorescent Dd strain. Additionally, the WT strains of both host and pathogen can easily be replaced with combinations of selected mutants, allowing to disentangle genetically whether the compound target is in the host or the pathogen. This is a way to guide the research towards functional genome wide screens, such as REMI-seq on Dd (Kuspa, 2006), Tnseq on Mm (Lefrançois et al., 2024), or dual RNAseq on both (Hanna et al., 2019). We are convinced that such a workflow will enable us to decipher mechanisms of cell autonomous defence pathways as well as virulence strategies with enough detail to boost or disarm them, respectively, eventually giving rise to new therapies to cure mycobacterial infections. This methods and protocols article follows on a recent methods article that illustrated the experimental versatility of Dd with a focus on microscopy techniques (Mottet et al., 2021), and aims at demonstrating with great technical detail the use of the high throughput Dd-Mm platform to test chemical and natural compounds for their antibacterial and anti-infective activities. In parallel, the generated data analysis workflow represents a powerful and adaptable tool for evaluating anti-infective compounds in high-throughput assays.

## Results

### Data analysis of plate reader assays

Data obtained from the BioTek Synergy H1 plate reader from infection or broth assays are processed in analogous manner with custom scripts written in R (Ramakrishnan, 2004). First, every well is annotated with the name of the test condition, the respective test concentration, the biological replicate as a number and the corresponding vehicle control. Vehicle controls wells are flagged with the prefix “VC” and positive controls with the prefix “PC”. Then annotated raw data are submitted to R to generate summary data. Briefly, the script loops over well plates, which are saved as separate xlsx files, extracts RLU and RFU for each well and calculates the area under the curve (AUC) for each well. Subsequently, the median, the median absolute deviation (mad), the mean and the standard deviation of the AUC of all vehicle controls are calculated. Additionally, we compute the median of the first measurement across all conditions of a well plate is computed and, extrapolated over the entire time course of the experiment in order to calculate the AUC of this baseline. These two values are used to linearly scale test conditions between 0 (the baseline AUC) and 1 (the vehicle control AUC), yielding one summary value per well (see Fig. 1J). Additionally, the robust Z’-factor (Atmaramani et al., 2020) is calculated for each vehicle control on each plate by using the respective vehicle control and the positive control as a reference. A pdf report containing plots, such as the temperature over time or heatmaps sorted by test condition are helpful for troubleshooting and visualization. In a second step, biological replicates are integrated, by looping over the respective summary data, notably the scaled AUC, and calculate the median over all technical replicates in all biological replicates for each condition. This workflow of normalized AUC is applied to both readouts, RFU and RLU in infection, and RLU in broth, to obtain a comparable and quantitative measure, which we call “normalized residual growth” (NRG). This allows to establish a hard cut-off for fast hit classification on every readout. From experience and integrating observed scatter of test conditions, we deemed an NRG of 0.5 a robust cut-off (see Fig. 1J). The NRG can take negative values in the case of a growth curve with a negative slope, i.e. decreasing under the starting value and thus indicating cell or bacteria killing, leading to an AUC below the extrapolated baseline. However, the assay is intended as a growth inhibition assay and not a killing assay, with highest resolution between the two references. A pdf report including plots which compare the technical replicates per biological replicate in a heatmap or correlate the scaled AUC to the AUC are helpful for troubleshooting and visualization. In case of dose response curves (DRCs), plots with the log_10_ of the test concentration on the x-axis and the scaled AUC on the y-axis are helpful for a fast overview over tested conditions. The half inhibitory concentration (IC_50_) was calculated on the NRG with GraphPad Prism (Version 8.0.1) using a robust 4PL regression, constraining the top value to 1 and, if necessary, the bottom value to -2 (Sebaugh, 2011). A minimum inhibitory concentration (MIC) was determined as the lowest experimental concentration, for which the averaged normalized residual growth plus or minus one standard deviation overlapped with 0 (Magréault et al., 2022).

**Fig. 1.**
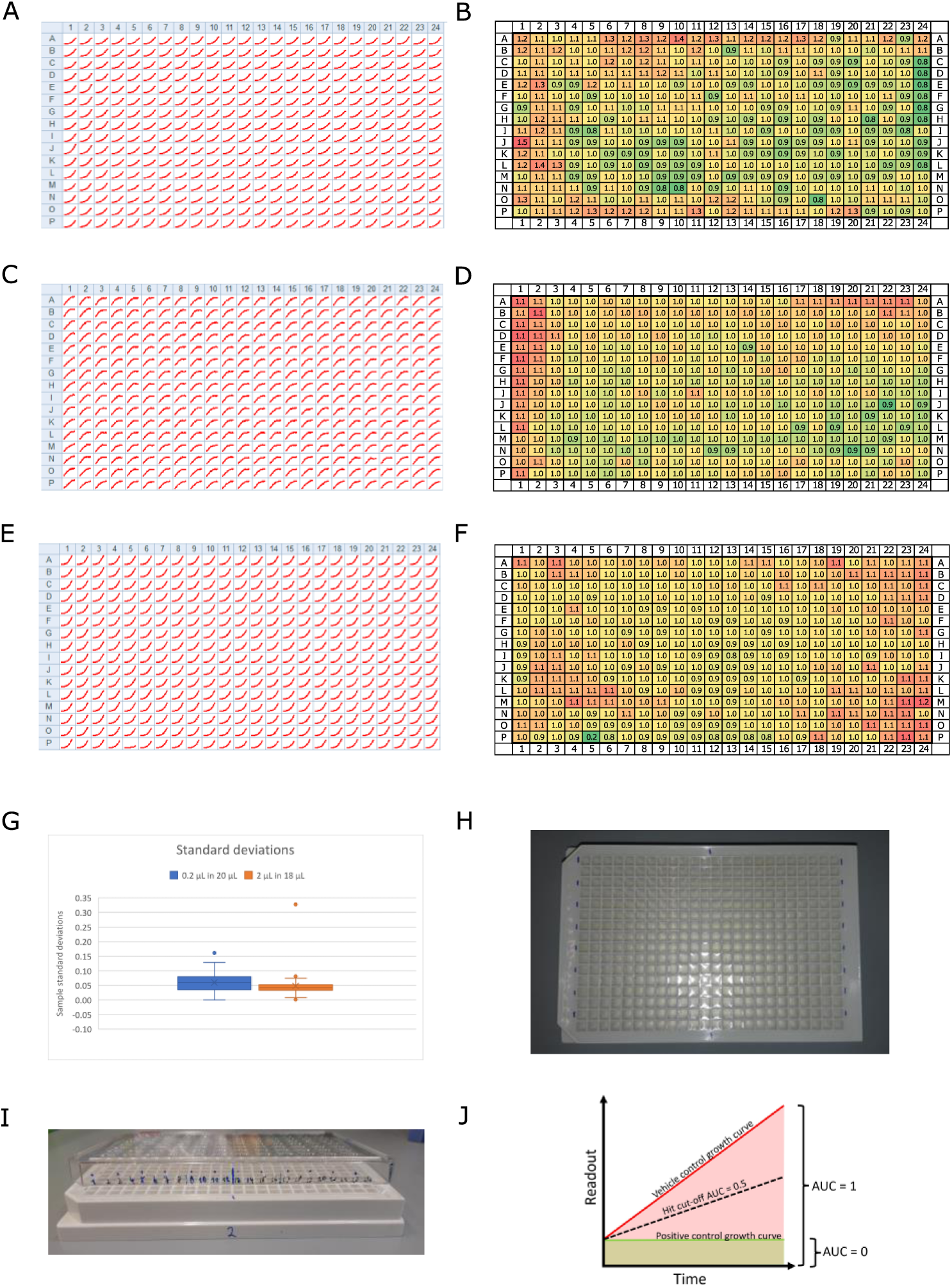
Overview on the assay design. Panel **A**, **C** and **E** show growth curves for every position on a 384 well plate for **A**, Mm in infection, **C**, Dd in infection, and **E**, Mm in broth. Homogeneity of these results across the full plate was assessed by dividing the maximum raw value of each well by the median of all maxima. The resulting rounded fold change is depicted in panels **B**, Mm in infection, **D**, Dd in infection, and **F**, Mm in broth. The color code indicates low values with green and high values with red tones. Panel G shows a box plot with the standard deviations of a set of about 50 compounds, which were tested at the same concentration on Mm in broth, once by addition in a 1:100 dilution (0.2 µL in 20 µL, blue) and once by addition in a 1:10 dilution (2 µL in 18 µL, orange). Panel **H** shows a photo illustrating labeling of the plate for easier pipetting. Panel I shows a photo of the same plate, with a labeled lid on top, illustrating a method to keep track of pipetted wells. Panel J illustrates the calculation of the normalized residual growth (NRG). The area under the curve of the vehicle control is scaled to 1, while the area under the curve of the positive control is scaled to 0.

### Benchmarking of the Dd-Mm high-throughput platform

The high-throughput *Dictyostelium discoideum*-*Mycobacterium marinum* (Dd-Mm) infection model was benchmarked with selected antibiotics commonly used in tuberculosis (TB) treatment and anti-infectives by generating dose-response curves (DRCs) and calculated IC_50_s and MICs where applicable (Fig. 2). The benchmarking involved two complementary assays. First, an anti-infective assay measured the intracellular growth of a bioluminescent Mm within Dd cells expressing mCherry. This setup also enabled simultaneous monitoring of the host amoebae’s growth. Second, an antibiotic assay assessed Mm growth in broth. In both assays, luminescence served as an indicator of bacterial biomass and metabolic activity. Bacteria or infected amoebae were plated in 384-well plates, treated with decreasing concentrations of each antibiotic, and monitored over 72 hours using a plate reader.

**Fig. 2.**
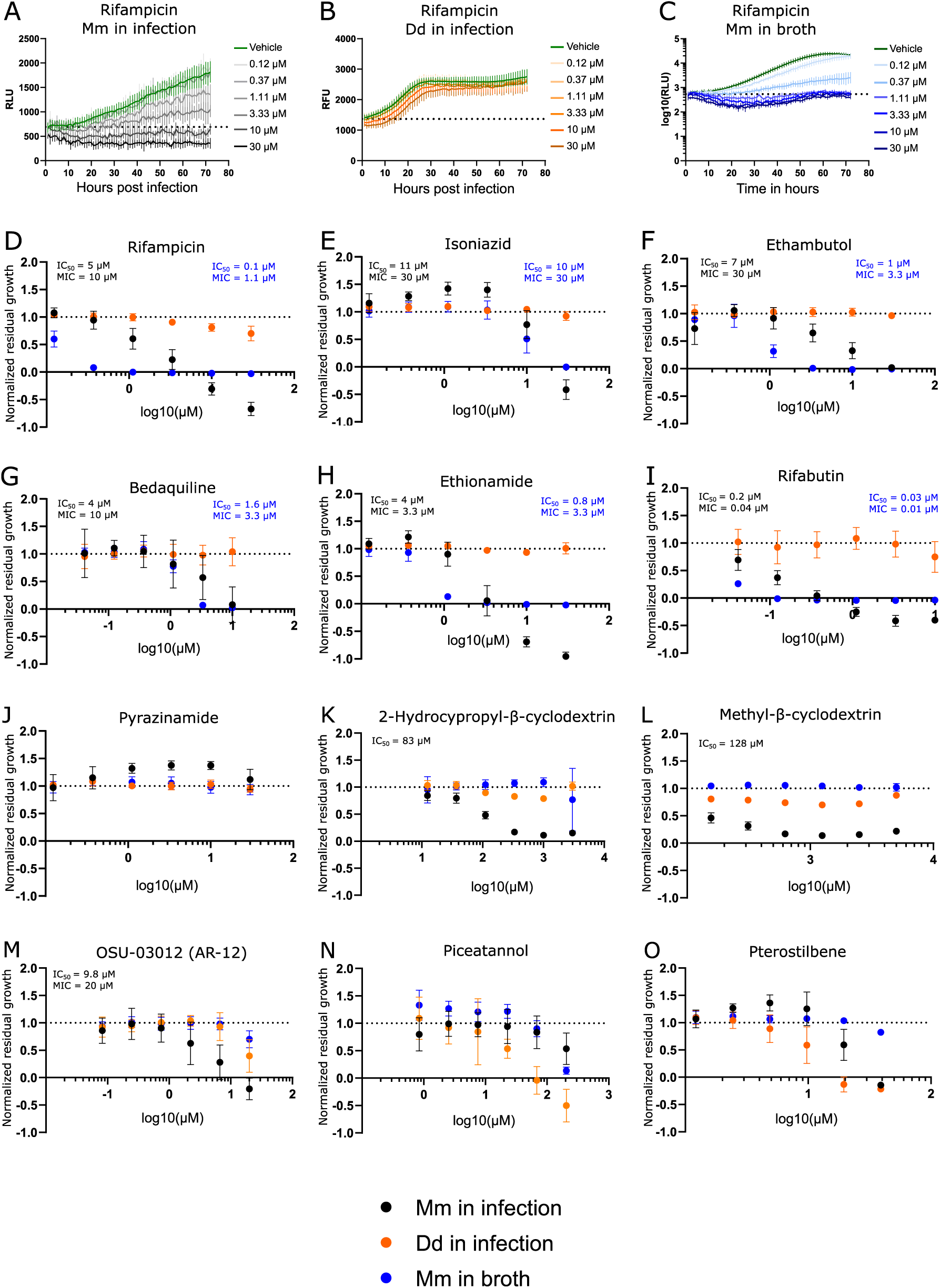
Dose response curves of benchmarked antibiotics, anti-infectives and Dd inhibitors. A selection of antibiotics used for treatment of a TB infection were benchmarked in the Dd-Mm high-throughput system, in infection and in broth. In panels **A**, **B** and **C**, 72 hour growth curves of Mm in infection, Dd in infection and Mm in broth, respectively, at different concentrations of rifampicin are shown. The dashed line represents the median over the first measurements of all wells of the respective experiment. On the y-axis are random luminescence units (RLU) or random fluorescence units (RFU). The former were log10 transformed in panel **C**. The corresponding dose response curve of Rifampicin is shown in panel **D**. The dose response curves depicted in panel **E** (isoniazid), **F** (ethambutol), **G** (bedaquiline), **H** (ethionamide), **I** (rifabutin) and **J** (pyrazinamide) were created from growth curves in analogous manner. Anti-infective compounds were benchmarked in panels **K** (2-hydroxypropyl-β-cyclodextrin), **L** (Methyl-β-cyclodextrin) and **M** (OSU-03012 or AR-12). Panels **N** and **O** show the stilbenes piceatannol and pterostilbene, respectively, both Dd growth inhibitors. Black points show Mm in infection, orange points Dd in infection and blue points Mm in broth. Log10 of the test concentration in µM is shown on the x-axis. A dashed line at y = 1 represents the normalized residual growth of the vehicle control. Depicted are means of at least three biological replicates with at least two technical replicates each and the respective standard deviations. Black and blue text inlays show the MIC and IC_50_ of the anti-infective or the antibacterial assay, respectively.

We observed susceptibility to first line antibiotics used against TB, rifampicin, ethambutol and isoniazid but not to pyrazinamide (see Fig.2 A-F, J). Indeed, Mm is reported to be naturally resistant to pyrazinamide (Aubry et al., 2000; Wagner & Young, 2004). In other susceptibility tests, it was also reported that Mm was susceptible to ethambutol and isoniazid only above certain breaking points (Aubry et al., 2000). Susceptibility to both antibiotics was observed, however notably for isoniazid at relatively high concentration in infection and in broth (MIC of 30 µM). Further, the susceptibility to bedaquilin, ethionamide and rifabutin, an analog of rifampicin was also detected (see Fig. 2G-I). The Dd-Mm system was also benchmarked with compounds having a higher activity in infection compared to in broth, compounds that we classify as “strict anti-infective”. For example, methyl-β-cyclodextrin and 2-hyroxypropyl-β-cyclodextrin reduced Mm growth in infection, but not in broth (see Fig. 2KL). In parallel, OSU-03012 (AR-12), a drug used as an autophagy inducer, was tested and seems to act as a strict anti-infective but in a specific and narrow dose range (Fig. 2M) (Chiu et al., 2009; Gao et al., 2008). We perturbed Dd growth with the two stilbenes piceatannol and pterostilbene and demonstrated that within a dose range Dd growth can be severely limited while still allowing for almost unperturbed Mm growth (see Fig. 2NO), indicating that Dd growth and intracellular Mm growth are partially independent. The average robust Z’-factor of the respective benchmarking experiments was 0.73 for Mm in infection and 0.61 for Mm in broth. Overall, we demonstrated the robustness of the Dd-Mm high through-put platform and its capability to differentiate between antibiotics and anti-infective compounds.

## Discussion

The data processing pipeline developed for the Dd-Mm infection and broth assays provides a robust and reproducible framework for analyzing high-throughput screening experiments. This workflow, implemented using custom R scripts, effectively automates data annotation, normalization, and summary generation, ensuring consistency and scalability in handling large datasets. Key steps in the pipeline include the annotation of test conditions, concentrations, and controls, followed by the calculation of area under the curve (AUC) for luminescence (RLU) and fluorescence (RFU) measurements. By scaling the AUC between the baseline and vehicle control values, the pipeline generates a normalized residual growth (NRG) metric, providing a standardized quantitative readout across plates. This normalization facilitates direct comparison between infection and broth assays and across biological replicates, enhancing the reliability of the results. The inclusion of robust statistical measures, such as the median absolute deviation (MAD) and the Z’-factor, ensures high-quality data. The Z’-factor calculation, using vehicle and positive controls as references, validates assay robustness and helps identify variability, a critical feature for high-throughput settings. Overall, this workflow represents a powerful and adaptable tool for evaluating anti-infective compounds in high-throughput assays. Its ability to integrate technical and biological replicates, normalize data across experimental conditions, and provide clear visualization supports confident decision-making in drug discovery. Future refinements could include expanding its application to killing assays or integrating machine learning algorithms for automated hit classification and outlier detection. The systematic and quantitative nature of this pipeline underscores its utility in high-throughput infection model platforms.

The susceptibility profile observed in the Dd-Mm infection model aligns well with known characteristics of Mm and its relationship to TB treatment. Susceptibility to first-line TB antibiotics rifampicin, ethambutol, and isoniazid was confirmed, with the exception of pyrazinamide confirming previous findings that Mm is naturally resistant to this drug The platform’s versatility was further highlighted by its ability to detect susceptibility to additional TB-relevant antibiotics, including bedaquiline, ethionamide, and rifabutin, a rifampicin analog. Notably, bedaquiline demonstrated not only its antibiotic activity but also potential host-beneficial effects, as previously reported (Giraud-Gatineau et al., 2020). These dual effects make it an attractive candidate for further investigation into host-pathogen dynamics. The benchmarking of compounds with higher activity in infection compared to broth—classified as “strict anti-infectives”—demonstrates the Dd-Mm system’s capability to distinguish between general antibiotics and compounds with infection-specific activity. For instance, methyl-β-cyclodextrin and 2-hydroxypropyl-β-cyclodextrin reduced Mm growth exclusively in the infection model, suggesting that these compounds may target host-pathogen interactions or intracellular bacterial survival mechanisms. Cyclodextrins have been reported as intrinsically bactericidal, which we did not observe at the tested doses. However, they are also speculated to manipulate the host-pathogen interface by depleting host membranes of sterol (Hammoud et al., 2019). Similarly, OSU-03012 (AR-12), an autophagy inducer, exhibited strict anti-infective activity within a narrow dose range, underscoring the platform’s sensitivity in detecting dose-dependent effects. Overall, the Dd-Mm system represents a powerful tool for investigating not only host-pathogen interactions but also the ability to distinguish between antibiotics and anti-infective compounds further underscoring its utility for drug discoveryand potentially identifying novel therapeutic strategies targeting intracellular pathogens.

## Material and methods

### 1. Preparation of media and cell culture material

#### 1.1. Mycobacterium marinum

##### 1.1.1. 7H9 for liquid cultures

- Use a measuring cylinder to dissolve 2.35 g of 7H9 powder (BD, Difco Middlebrook 7H9) in 450 mL double distilled water.
- Add 1 mL Glycerol (Sigma Aldrich 42025, suitable for cell culture, final 0.2 %)
- Add 250 µL of Tyloxapol (Merck T8761-50G, final 0.05 %). To reduce the viscosity of Tyloxapol and make it easier to pipette, heat it briefly in a microwave for easier pipetting.
- Stir until the solution is clear.
- Filter-sterilize (Steritop, 0.22 µm PES) under a laminar flow hood, and add 50 mL of OADC (BD BBL, Middlebrook Oleic Albumin Dextrose Catalase Growth Supplement, aliquots filter-sterilized beforehand and stored at 4 °C).
- Store medium at 4°C until use. Prewarming is not necessary

##### 1.1.2. 7H11 for agar plates

- Dissolve 11.67 g of 7H11 Agar (BD, Difco 7H11) in 500 mL double distilled water, directly in a 500 mL bottle.
- Add 2.5 mL Glycerol (final 0.5 %), mix thoroughly by shaking and autoclave. Autoclaving will dissolve agar completely.
- Use immediately after autoclaving or reheat later using a microwave. When the agar is liquid after autoclaving or microwaving, take the necessary volume of agar to plate.
- Add OADC (10 % of the total volume)
- Add the appropriate antibiotic selection (Kanamycin at 50 µg/mL, hygromycin at 100 µg/mL, rifabutin at 10 µg/mL final concentration) and plate approximately 20 mL per 10 cm Petri dish. Leave the Petri dishes standing for at least 30 min under the hood to cool and dry.
- Stored at 4 °C, plates can be used up to 1 month.

#### 1.2. Dictyostelium discoideum

##### 1.2.1. Filter-sterilized HL5-C medium

- To prepare five bottles of 900 mL each, dissolve 119.25 g of HL5-C powder (Formedium, powder to be stored at 4°C) in 4.5 L of double distilled water. Shake the powder bottle before use to ensure homogenous medium preparations.
- Stir for at least 1 h until fully dissolved.
- Check pH with pH paper, pH = 6.5.
- Filter-sterilize under a laminar flow hood into 1 L Schott bottles and store at 4 °C up to 3 months until further use. Prewarming to room temperature is recommended. After opening a bottle for the first time, store at room temperature.

##### 1.2.2. Antibiotics in the culture medium

Alternatively, Penicillin/Streptomycin (P/S, 10 mg/mL and 10 000 U/mL, aliquots at -20°C, final concentration in medium 100 µg/mL and 100 U/mL) can be added once a bottle is used first. This prevents contaminations but can make it harder to detect resistant contaminations. It is not recommended to add P/S in the maintenance culture.

##### 1.2.3. Fluorescence of HL5-C in GFP channel

HL5-C has a significant autofluorescence background at GFP emission wavelengths. This is problematic, since it makes segmentation of images acquired at the high content microscope difficult and adds an offset signal to readouts acquired with a plate reader. This background signal could not be attributed to a specific ingredient in the medium, but storage at room temperature under constant lighting decreased it.

## 2. Maintenance of cell and bacteria cultures

### 2.1. Mycobacterium marinum

#### 2.1.1. Cycle of plate culture, primary culture, and liquid stocks

A fresh culture of Mm LuxCDABE (Andreu et al., 2010) or Mm pMSP12::GFP (Addgene plasmid # 30167) (Davis et al., 2002) (derived from *M. marinum* M strain ATCC BAA-535) is plated every three months. Starter cultures and liquid stocks from a plate culture should be used maximum one month before preparing a new starter culture. Generally, passage numbers should be kept low for mycobacteria culture and it is recommended to use a primary liquid culture as an inoculum for each infection experiment.

#### 2.1.2. Plate culture and primary liquid culture

- Recover Mm from a -80 °C glycerol stock and start a plate culture using an inoculation loop then streak onto the 7H10 agar plate.
- After sealing the plates with parafilm, incubate at 32 °C in a box, maintain humidity by adding a wet paper tissue. After three days, growth should be visible, depending on the size of the inoculum, after ca seven days the colonies should have grown sufficiently to store the plates at 15 °C.
- Start a primary liquid culture (10-20 mL) from the 7H10 plate culture in an Erlenmeyer flask with a gas permeable plug.
- Syringe the inoculum ten times using a blunt needle (25 Gx3/4” blunt needles, preferably 3 mL syringes, aspirating and releasing once means the suspension has been syringed once). Make sure to use a needle-proof glove to prevent injury.
- Prepare the Erlenmeyer flask with 7H9 medium and antibiotic selection (see Supplementary table 3)
- Add the syringed inoculum to the Erlenmeyer flask.
- Incubate the culture at 32 °C, shaking (150 rpm) for at least 24 hours.
- This culture can now be used to inoculate liquid cultures for infection experiments, while the rest can be stored in the fridge at 15 °C. Liquid stocks should be used for maximum one month.

##### No glass beads in liquid cultures

Note: Glass beads of 3 mm diameter can be used in liquid cultures to limit clumping by increasing shear stress. However, alternative methods such as the use of Tyloxapol instead of Tween80 or a short centrifugation of 20 g for 1 min seems to be sufficient to eliminate Mm clumps.

##### Measuring OD with the Ledetect 96

Use the plate reader to read OD in transparent 96 well plates. Use 200 µL of bacterial suspension per well and make sure no bubbles perturb the light path in the well. Measure OD at 620 nm and multiply the result by 2 to obtain the pathlength corrected OD. Although 600 nm are the traditional wavelength to measure OD, the machine’s closest wavelength is 620 nm. The difference in OD at 600 and 620 nm was examined with a photometer and found negligible. The pathlength correction extrapolates the OD obtained across a 96 well plate well to the pathlength of a photometer cuvette (1 cm).

### 2.2. Dictyostelium discoideum

- Recover Dd Ax2(Ka) expressing mCherry from frozen spore stocks (SoerensenMC with 10 % glycerol, see Supplementary table 3) stored in liquid nitrogen. A culture should not be used for infection before having been passaged for one week and showing normal doubling times. Infection efficiency decreases with the age of a culture, after one month a fresh culture needs to be started as described above. It is recommended to keep record of the passage number of the current culture.
- Dd Ax2(Ka) maintenance culture should be performed following a standard protocol (Bloomfield et al., 2008; Fey et al., 2007). Briefly, cells are cultivated in adherent conditions in HL5-C medium at 22 °C under constant light to a maximum density of 5*10^7^ cells/10 cm Petri dish since overgrowing has been observed to result in low infection efficiency.
- We use a splitting regime as follows: Monday (1/10 dilution), Wednesday (1/10 dilution), Friday (1/20 – 1/50 dilution).

#### Measuring cell density with the Countess

Countess II FL (life technologies, slides by Invitrogen) is used to assess cell density. After pipetting 10 µL of cell suspension into the measurement slide, incubate 1 min to allow cells to sediment in the focal plane before measuring by inserting the slide and select only the brightfield channel. The resulting cell count must be divided by 2 since the device assumes a 1:2 dilution in trypan blue and automatically corrects for it. Take at least two measurements to exclude measurement fluctuations. If necessary, adjust the focus manually. The practical measuring range was reported to go up to ca 6*10^7^ cells/mL.

## 3 Preparations for the infection

### 3.1. Mycobacterium marinum

- From the primary liquid culture, inoculate a liquid culture for an infection. The culture should be started ca 24 hours before infection.
- For the infection culture, use an Erlenmeyer flask with a gas-permeable plug. After adding an appropriate volume of 7H9 (between 10 and 30 mL of final culture volume) and the appropriate antibiotic, prepare the inoculum from the stored primary liquid culture.
- Resuspend by vortexing and subsequently pellet clumps by spinning at 300 rpm (corresponds to 20 g with the Thermo Scientific™ 75003180 rotor) for 1 min.
- Take 2-3 mL of the supernatant and syringe (as described above) before adding the inoculum to the Erlenmeyer flask.
- Incubate the culture at 32 °C, shaking (150 rpm). Measure OD (as described above) to ensure adequate growth for the following day. If the culture is prepared 24 h before the experiment, typically a pathlength corrected OD of ca 0.3 is sufficient to obtain an OD = 1 on the day of the experiment.

### 3.2. Dictyostelium discoideum

- Plate Dd mCherry cells 24 hours prior to the experiment in 10 cm Petri dishes at 1*10^7^ cells per Petri dish (around 40% of confluency), in filter-sterilized HL5-C medium without penicillin/streptomycin or additional selection to avoid residual antibiotic compromising bacterial virulence during infection.
- Incubate for 24 hours under normal growth conditions at 22 °C. The plates should be approximately at 3-4*10^6^ cells per mL (3-4*10^7^ cells per Petri dish), corresponding to 90-95% of confluency on the day of infection. The adherent cell lawn will be infected by spinoculation, therefore it should be as dense as possible while the number of floating cells should be minimal.

## 4. Infection

- Decant the Mm culture from the Erlenmeyer flask into a 50 mL conical tube (Falcon) and spin down (20 g for 1 min, as described before).
- Aspirate the supernatant and measure the OD (as described before). From the OD, calculate the bacteria density (see calibration curve of OD to bacteria in supplementary figure 1, supplementary table 1) and transfer the volume that corresponds to 8.75*10^8^ bacteria (MOI of 25 assuming 3.5*10^7^ Dd per Petri dish) to a fresh conical tube. During subsequent steps, material loss might happen, consequently it is recommended to include an excess of 20%, e. g. 1.2 * 8.75*10^8^ bacteria.
- Centrifuge the mycobacteria at 2,700 g for 10 min (corresponds to 3,500 rpm with the Thermo Scientific 75003180 rotor). After centrifugation, discard the supernatant by decanting and remove leftovers of 7H9 subsequently by pipetting carefully.
- During the centrifugation step, aspirate the medium from the Dd culture and add 5 mL of filter-sterilized HL5-C medium without penicillin/streptomycin or selection.
- Resuspend in 600 µL (500 µL plus a margin of 20% = 600 µL) of HL5-C per Petri dish to be infected.
- Syringe the suspension ten times through a blunt needle to break up clumps, as described above for the preparation of the infection culture. The suspension should be syringed 10 times at the maximum, to avoid damage to the bacteria. Bubbles should also be avoided. It is good practice to visually check the suspension under the microscope to ensure the effectiveness of syringing.
- Add 500 µL of the bacterial suspension to each Petri dish.
- Shake the Petri dish gently crosswise. Then seal the Petri dish with parafilm.
- Centrifuge twice at 500 g for 10 min (corresponds to 1,500 rpm with the Thermo Scientific™ 75003180 rotor) to accelerate sedimentation of bacteria and therefore increase contact between Dd and Mm Turn the bucket by 180° between spins to redistribute the bacteria suspension. Shake the Petri dishes in the bucket crosswise between the first and second spin.
- Incubate for 10-20 min to allow phagocytosis. This time is not very critical but should stay coherent throughout experiments. In case of a timeline experiment, the start of the phagocytosis step marks 0 hpi (hours post infection).
- Wash off the extracellular bacteria with 7-10 mL of HL5-C several times (3 – 8 repeats, depending on the subsequent experiment. For instance, five repeats are recommended for a plate reader experiment). The washes should result in as few extracellular bacteria as possible, to avoid a false positive signal. All Petri dishes should be treated equally. Special attention should be paid by adding and removing the medium on the edge of the Petri dish to avoid detaching Dd cells.
- After washing, cells are detached mechanically, resuspended in HL5-C containing fresh 5 µg/mL and 5 U/mL P/S (1/2000 dilution of a 10 000 µg/mL stock or 25 µL of P/S in 50 mL HL5-C, see note on use of P/S) and kept in a conical tube for further use.

### Multiplicity of infection (MOI)

The MOI depends on the number of Dd and the number of Mm. The former is not directly quantifiable since the cells must remain adherent for the infection but is instead estimated in the range from 3-4*10^7^ cells per 80-90% confluent Petri dish from previous experiments, combined with visual inspection of the confluency.

### Note for *Mm* ΔRD1mutant

Some Mm mutant are attenuated during infection and may require an adaptation of the MOI such as the ΔRD1 mutant for which the MOI is doubled in order to compensate for its lesser phagocytosis. Due to the higher MOI used for such mutants, additional washes are necessary to eliminate extracellular bacteria.

## 5. Monitoring the course of infection

### 5.2. Intracellular growth of *Mycobacterium marinum* during infection

- To prepare cells for a growth assay in a 96 (luminescence: Thermo Scientific 136101) or 384 well plate (luminescence: Interchim FP-BA8240) using Mm LuxCDABE (Andreu et al., 2010) or Mm pMSP12::GFP (Addgene plasmid # 30167) (Davis et al., 2002), measure cell density at the Countess and prepare the final cell suspension via a 1:10 intermediate dilution.
- In 96 well plates, 2.5*10^5^ cells/mL (5*10^4^ cells per well in 200 µL) are seeded.
- In 384 opaque-bottomed well plates (Interchim FP-BA8240) 5*10^4^ cells/mL (10^3^ cells per well in 20 µL) are seeded.
- Prepare a cell suspension (ca 20 mL per 96 well plate, ca 10 mL per 384 well plate) at the correct density in a conical tube. Cells can be plated using a multipipette (Sartorius Picus, 12 channel, 5-120 µL) or an automatic dispenser.
- In the case of 96 well plates, it is recommended to fill the interwell space with 150 µL of double distilled sterile water minimizing the “edge effect” of extreme growth at the edge of the well plate (see Fig. 1A-F).
- It is recommended to briefly spin down 384 well plates to ensure the liquid is at the bottom of the well, both after plating cells and after adding test compounds.
- Add test compounds in a 1:100 dilution: 0.2 µL in 20 µL, ideally using an electronic multipipette (see “Note on compound addition”, Fig. 1G).
- Seal plates with a gas-impermeable (ROTILABO, H769.1; 384 WPs) or gas permeable seal (4titude, 4ti-0516/96; 96 WPs). Since the airtight seals come non-sterile, it is recommended to sterilize them using the UV program of one of the laminar flow benches.
- To limit the edge effect of stacked plates it is recommended to place an empty, unsealed dummy plate at the first and last positions of the stack.
- Luminescence and fluorescence are read from the top, with gain values signal saturation (ca gain = 100 for both readout with a BioTek Synergy H1 reader), and an integration time of 1s (higher integration time does not impact RLU or RFU but decreases the noise of the measurement. If time resolution is not a concern, 10s of integration time is recommended).

#### Note on using a stacker and controlling temperature

When using the BioStack, the assay plates spend most of the 72 hours assay in the stacker. Consequently, the ambient temperature should be controlled to ensure growth of bacteria and amoebae. Both organisms have different temperature optima, and the optimal compromise is at 25°C. However, the BioTek Synergy H1 plate reader is placed in the same environment as the BioStack and can heat, but not cool the space around the plate carrier. Consequently, the inside of the plate reader will reach temperatures of around 30°C. To avoid such a temperature shock to the amoebae, we set the ambient temperature to 24°C, allowing for growth while the samples are in the stacker and minimizing a heat shock while they are measured.

#### Note on compound addition

Different strategies were tested for optimal addition of the compounds to 384 well plates: 1. A dispenser (Agilent Bravo liquid handler) in a 1:10 dilution, 2. an electronic multipipette (Sartorius Picus, 12 channel, 0.2-10µL) in a 1:10 dilution and 3. a 1:100 dilution and 4. a single channel pipette (Gilson, P2) used with a 1:100 dilution. Options 2 and 3, i.e. using an electronic multipipette with a 1:100 or a 1:10 dilution (i.e. adding 0.2 µL in 20µL or 2 µL in 18µL, respectively) was better, due to good control of the added volume. Notably, we observed a negligible difference in measurement scatter between using a 1:10 or a 1:100 dilution (Fig. 1G), consequently we preferred a 1:100 pipetting regime, since it gave us more flexibility in handling stocks of test conditions. Additionally, we recommend a comprehensive well plate design, for example we used sector four (i. e. every bottom right well of a quadruple well) exclusively for controls. Generally, vehicle controls were homogenously distributed over the plate, with the positive control (10 µM rifabutin) in the top and the bottom rows. At least three biological replicates were acquired (i. e. different Petri dishes of infected amoebae) with 3, 2 or 1 technical replicate each, depending on availability of the extracts or compounds.

### 5.3. Growth of *Mycobacterium marinum* LuxCDABE in broth

- Prepare the Mm culture as described for the infection.
- On the experiment day, directly take 1 mL of the culture supernatant (after centrifuging for 1 min at 300 rpm, as for the infection) and transfer to an Eppendorf tube.
- Syringe as described in the infection protocol.
- Measure the OD of the syringed culture supernatant as described before.
- Calculate the required volume of culture supernatant to prepare a suspension at 3.75*10^5^ bacteria/mL (7,500 bacteria per well in 20 µL).
- Prepare this suspension via a 1:10 dilution of the syringed culture supernatant (900 µL 7H9 + 100 µL culture supernatant) to reduce the margin of error due to pipetting small volumes..
- Fill 384 well plates with an electronic multipipette and centrifuge briefly before adding test extracts, compounds or mixtures.
- For the use of the stacker or plate reader, proceed as described above for the infection assay.

#### Note on using a stacker and controlling temperature

As described above, the fact that the plate reader cannot cool its internal temperature, while being placed in a heated environment, can lead to a heat shock for the samples. In case of the “in broth” assay, we set the ambient temperature to 27°C, leading to ca 30°C inside the plate reader. These temperatures are indeed below the optimal growth temperature of Mm but were observed to dampen an edge effect through sacrificing overall growth.

## Supplementary information

### 1. Generation of *M. marinum* stocks

- Glycerol stocks of Mm are generated from a liquid culture by adjusting the culture medium to 20% glycerol and aliquoting the desired amount into tubes.
- The stocks can then directly be stored at -80°C or in liquid nitrogen.

### 2. Generation of *D. discoideum* spore stocks

- Grow Dd in suspension (HL5-C medium, 22°C, shaking at 120 rpm) until a density of 1*107 cells/mL.
- Pellet the cells in SoerensenMC by centrifugation (500 g, 10 min, corresponds to 1500 rpm with the Thermo Scientific™ 75003180 rotor), add back SoerensenMC to repeat the wash to eliminate traces of HL5-C.
- Prepare agar dishes with SoerensenMC and 2% Bacto agar (BD, Difco Bacto Agar).
- Plate 2 mL of cell suspension per agar plate, if necessary, remove excess liquid by pipette after allowing cells to sediment on the agar plate.
- Put lids on and incubate upside down at 22°C for at least 24h (up to 3 days) in a closed box with humid paper towels to prevent drying out of the plates.
- After fruiting bodies are generated, smash them on the agar by tapping the Petri dish vertically. Then, wash spores from the plate with SoerensenMC. For this, add 1 mL of SoerensenMC, swirl, and collect until most material is removed from the agar dish. Pool the suspension from multiple Petri dishes, if necessary.
- Measure spore density with the Countess (as described above for cell suspensions).
- Wash the spore suspension by centrifuging (500g, 10 min, corresponds to 1500 rpm with the Thermo Scientific™ 75003180 rotor), decanting the supernatant and resuspending in a volume of 10% glycerol in SoerensenMC to adjust the density to ca 107 spores per mL.
- Freeze in aliquots from 0.2 to 1mL and store overnight at -80°C in a Nalgene freezing box with isopropanol before transferring tubes to storage at -80°C or in liquid nitrogen.
- Check quality of the stock by thawing one aliquot and passaging the cells for one week.
- SoerensenMC 10x:
- Dissolve KH2PO4 19.97 mg; Na2HPO4*H2O 3.56 mg; MgCl*6H2O 0.1 mg; CaCl2*6H2O 0.1 mg in 1L double distilled water.
- Filter-sterilize, store at room temperature, dilute 1:10 in double distilled water for SoerensenMC 1x, adjust pH to 6.0 with NaOH or H3PO4.

### 3. *M. marinum* quantification

For calibration of optical density measurement to density of a liquid Mm culture, we used the optical bacterial cell counter Quantom Tx^TM^ (Logos Biosystems). Briefly, we grew bioluminescent or GFP expressing Mm to ODs up to 1.8 prepared serial dilutions and measured the respective OD. Subsequently we prepared the samples according to the manufacturers recommendations to be analyzed with the cell counter. Correlating the results with the previously measured ODs yielded the calibration curve presented and a linear fit resulted in the following formula for the bacterial density: 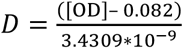

**Supplementary fig. 1:**
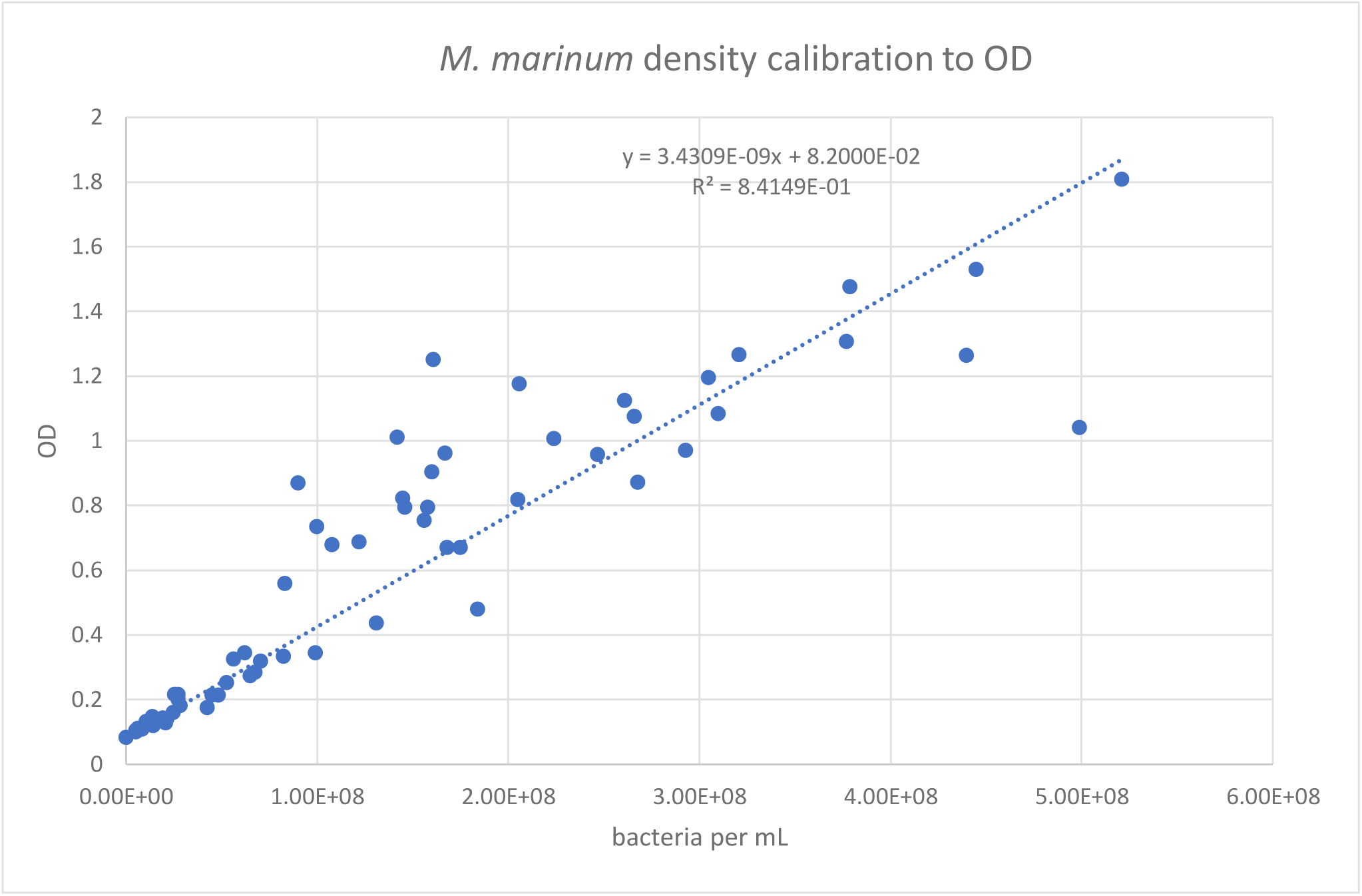
OD calibration curve. The x-axis shows bacteria per mL as measured by the Quantom Tx^TM^, the y-axis shows optical density of the sample. Each data point represents one measured sample. The curve pools samples from two strains, a strain expressing the bacterial *lux*-operon and a strain expressing GFP.

### Supplementary table

Correspondence table for the OD-bacterial density calibration curve. The left column shows the interpolated OD equivalents of the interpolated bacterial density in the right column.

| OD | bacteria/mL |
| --- | --- |
| 0.4 | 9.27E+07 |
| 0.45 | 1.07E+08 |
| 0.5 | 1.22E+08 |
| 0.55 | 1.36E+08 |
| 0.6 | 1.51E+08 |
| 0.65 | 1.66E+08 |
| 0.7 | 1.80E+08 |
| 0.75 | 1.95E+08 |
| 0.8 | 2.09E+08 |
| 0.85 | 2.24E+08 |
| 0.9 | 2.38E+08 |
| 0.95 | 2.53E+08 |
| 1 | 2.68E+08 |
| 1.05 | 2.82E+08 |
| 1.1 | 2.97E+08 |
| 1.15 | 3.11E+08 |
| 1.2 | 3.26E+08 |
| 1.25 | 3.40E+08 |
| 1.3 | 3.55E+08 |
| 1.35 | 3.70E+08 |
| 1.4 | 3.84E+08 |
| 1.45 | 3.99E+08 |
| 1.5 | 4.13E+08 |

### 4. Calibration of RFU to *D. discoideum* cell number in the plate reader

The calibration of bioluminescent Mm to RLU obtained in a plate reader, has already been documented (Andreu et al., 2010; Trofimov et al., 2018). To demonstrate the linear relationship between cell density and RFU measures in a plate reader monitoring of Dd expressing mCherry from the *act5*-locus (Paschke et al., 2019), cultures were grown in suspension to a high density, performed a serial dilution, measured the samples with the Countess (as described above) in four technical replicates, plated 20 µL of the same sample in four technical replicates into a 384-well plate (Interchim FP-BA8240) and obtained RFU under the same parameters as for our growth assay.

**Supplementary fig. 2:**
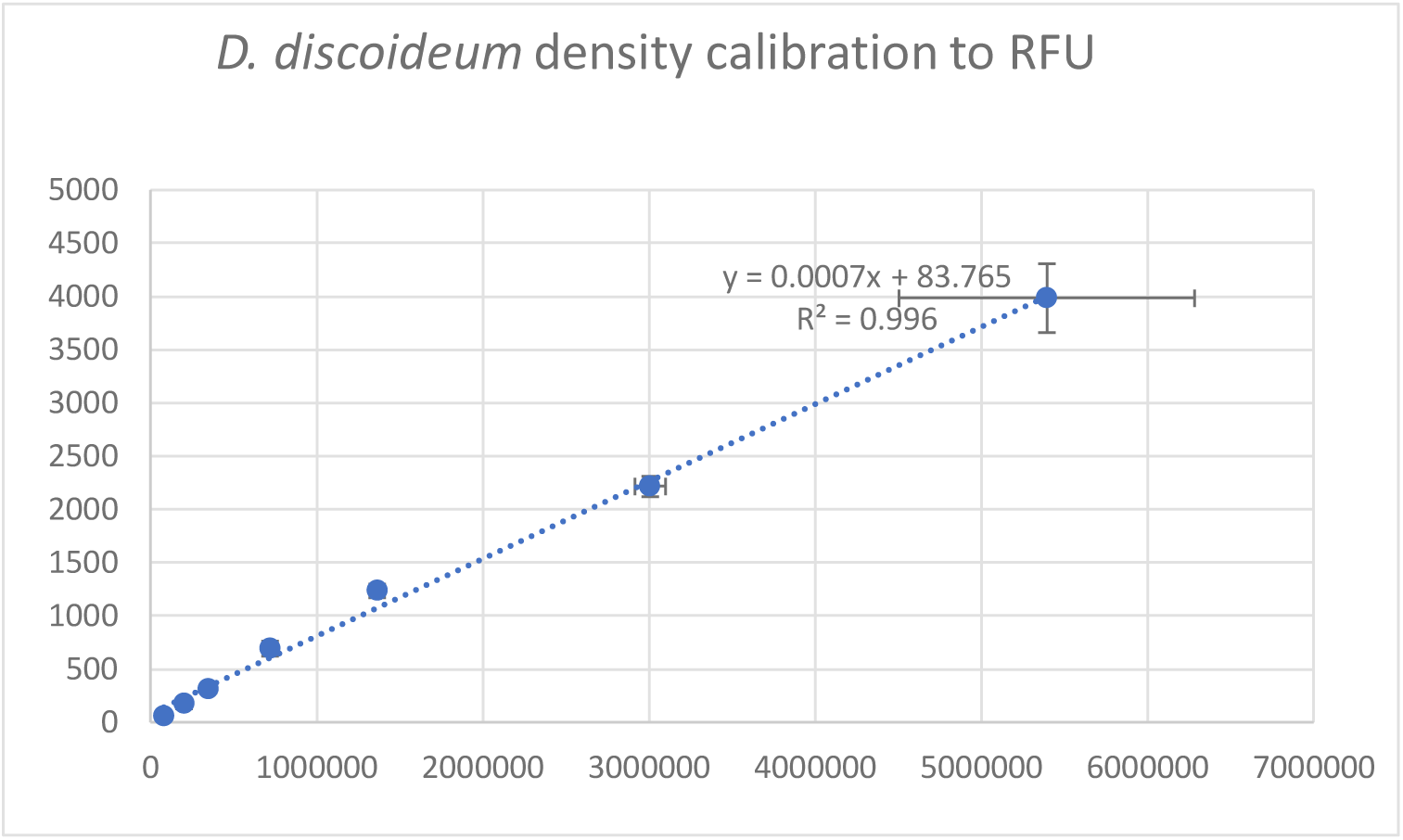
RFU calibration curve. The x-axis shows averaged cells per mL as measured by the Countess cell counter, the y-axis shows averaged RFU of the sample. Each data point represents the average over four technical replicates of measurements from both readouts. The error bars are the corresponding standard deviations.

### 5. Overview of used multi-well plates

**Supplementary table 1:** Overview over used well plates and their measures. This table gives an overview over well plates, their capacity, reference, provider, shape of their wells, diameter or edge length and resulting bottom surface area per well.

| Well plate and capacity | Reference number and manufacturer | Round or square | diameter/edge length in mm | bottom area in cm<sup>2</sup> |
| --- | --- | --- | --- | --- |
| 96 WP opaque | Thermo Fischer 136101 | Round | 6.55 | 0.34 |
| 384 WP opaque | Interchim FP-BA8240 | Square | 3.65 | 0.13 |

### 6. Overview over strains and constructs

**Supplementary table 2:** Overview over used strains and constructs.

| Strain | Construct name | Selection | Addgene # | Reference |
| --- | --- | --- | --- | --- |
| *D. discoideum* mCherry | pDM1514 | Hygromycin 50 µg/mL | 108999 | (Paschke et al., 2019) |
| *M. marinum* LuxCDABE | pMV306hsp | Kanamycin 50 µg/mL | 26155 | (Andreu et al., 2010) |
| *M. marinum* GFP | pMSP12::GFP | Kanamycin 50 µg/mL | 30167 | (Davis et al., 2002) |

